# Off-target genomic effects in MRP8-Cre driver mice complicate its use in weight gain and metabolic studies

**DOI:** 10.1101/2025.01.06.631557

**Authors:** Fatemeh Soltani, Cedric Duval, Robert A. S. Ariëns, Mari T. Kaartinen

**Affiliations:** Faculty of Medicine and Health Sciences (Division of Experimental Medicine), McGill University, Montreal, QC, Canada; Leeds Thrombosis Collective, Discovery & Translational Science Department, Leeds Institute of Cardiovascular and Metabolic Medicine, University of Leeds, Leeds LS2 9NL, United Kingdom; Faculty of Dental Medicine and Oral Health Sciences (Biomedical Sciences), McGill University, Montreal, QC, Canada

**Keywords:** Obesity, Adipose tissue, Immune system, Thrombosis, Factor XIII-A, MRP8-Cre model

## Abstract

**Background:** Thromboinflammation of adipose tissue involves accumulation of pro-fibrinogenic factors to adipose tissue in obesity, which promotes immune cell infiltration, affects weight gain and can lead to metabolic dysfunction. The role of neutrophil genes in inflammation are frequently investigated using MRP8-Cre mice to generate neutrophil-specific knockouts. Recent study demonstrated that MRP8-Cre mice have off-target deletions in *Serpine1* and *Ap1s1* genes. *Serpine1*, encoding plasminogen activator inhibitor-1, is a key anti-fibrinolytic factor linked to thromboinflammation and metabolic dysfunctions in obesity. In this study, we provide evidence suggesting a critical limitation in using MRP8-Cre model to study adipose tissue, weight-related dysfunctions and/or metabolic disorders.

**Methods:** MRP8-Cre and *F13a1*^-/-MRP8^ mice were placed on either control diet or high-fat diet for 16 weeks, with body weight monitored weekly. The expression of *Serpine 1, Ap1s1* and *Adgre1* (macrophage marker) genes in inguinal and epididymal adipose tissues were analyzed using qRT-PCR and compared to the wild-type mice.

**Results:** MRP8-Cre shows no *Serpine1* or *Ap1s1* expression in inguinal and epididymal adipose tissues. MRP8-Cre mouse is resistant to weight gain on obesogenic diet and does not show macrophage marker in adipose tissue compared to control obesity model. The resistance to weight gain translates to a neutrophil knockout model, *F13a1*^-/-MRP8^ that was not expected to show resistance to weight gain as its global knockout does not exhibit this phenotype.

**Conclusion:** Our work suggests that MRP8-Cre model may not be suitable to investigate metabolic outcomes of neutrophil genes, or pathologies that have underlying etiology in thomboinflammation.

## 1. Introduction

Obesity is a major risk factor for cancers and several metabolic and thrombotic diseases [1-3]. Several obesity comorbidities have underlining contributions from the chronic, systemic low-grade inflammation, which originates from adipose tissue (AT) dysfunction during weight gain [4-6]. Adipose tissue inflammation arises from several factors, including adipocyte stress and death, mechanical stress on cells from surrounding fibrotic extracellular matrix (ECM), and dysregulation of fatty acid metabolism [7-9]. The inflammation hampers with adipogenesis and lipogenesis which in turn causes ectopic lipid accumulation to the metabolic organs, such as liver and skeletal muscle [10, 11].

AT inflammation is initially driven by neutrophils whose activation precedes the pro-inflammatory macrophage infiltration. Macrophage arrival to AT is promoted by the presence of fibrin deposits and pro-coagulatory and anti-fibrinolytic molecules such as thrombin, Factor XIII-A (FXIII-A) transglutaminase, and plasminogen activator inhibitor-1 (PAI-1) in AT [12-14]. The process, referred to as thromboinflammation, is crucial in obesity-related metabolic dysfunction. Knocking out PAI-1, a key anti-fibrinolytic factor, in mice leads to resistance to weight gain and protection from AT inflammation and obesity-related metabolic dysfunctions [15-19]. Similarly, thrombin inhibitors administered during high fat diet (HFD) decrease weight gain [12] and transgenic mice with the mutation in fibrinogen that eliminates leukocyte integrin binding site, exhibit decreased AT inflammation and resistance to weight gain [12]. The global knockout of fibrin crosslinking enzyme, FXIII-A on a HFD displays normal weight gain but improved insulin sensitivity, decreased ECM in AT, and reduced macrophage infiltration[13].

Neutrophils play a significant role in obesity and thrombotic disorders [20-23]. Upon weight gain, neutrophils infiltrate adipose tissue within days, promoting pro-inflammatory macrophage recruitment and thereby worsening adipose tissue inflammation [24]. Neutrophils contribute to this through mechanisms such as secreting inflammatory cytokines and releasing neutrophil extracellular traps (NETs) which contribute to pro-thrombotic state and thromboinflammation [25, 26]. As first responders to injury sites, they also contribute to hemostasis and thrombus formation [27-30]. Research models and methods to study the effects of neutrophils on whole body biology include pharmacological depletion of neutrophils [31] and neutrophil specific knockouts generated using LysM-Cre and MRP8-Cre-ires/GFP (MRP8-Cre) mice. LysM-Cre is not specific to neutrophils, making the MRP8-Cre model more suited in this specific context [31-33]. MRP8-Cre-model expresses Cre recombinase via MRP8 (S100A8) promoter [34-36]. A recent study reported off-target effects in MRP8-Cre [31]. Authors reported that integration of a single copy of MRP8-Cre-ires/GFP transgene into chromosome 5 leads to the complete deletion of *Serpine1 (*PAI-1) gene and partial deletion of *Ap1s1* (Adaptor Related Protein Complex 1 Subunit Sigma 1) genes in the host genome [37]. A recent report [38] highlighted a critical limitation of the MRP8-Cre model showing that the MRP8-Cre/ires-eGFP transgene predisposes mice to cartilage antibody-induced arthritis. The severity of arthritis was attributed to the MRP8-Cre model itself, of which neutrophils showed compromised PAI-1 expression [38]. Herein, we present further evidence that MRP8-Cre may not be suitable for studies due to the off-target deletions. We show that MRP8-Cre subcutaneous (inguinal) and visceral (epididymal) AT do not express *Serpine 1* mRNA, and the mice phenocopy resistance to the published lack of weight gain of the global PAI-1 knockout on HFD. We further support this claim by showing that *F13a1*^-/-MRP8^ model created with MRP8-Cre mice also exhibits resistance to weight gain upon HFD, which is a phenotype not found in published study on global *F13a1* null mice [13]. Our finding suggests the careful evaluation of this Cre-model also in weight gain and metabolic studies and including it as an important control to any studies where PAI-1 and AP1S1 may have functions.

## 2. Materials and methods

### 2.1. Animals and MRP8-Cre

All animal housing and procedures were conducted according to the guidelines for animal experimentation and were approved by the Animal Care Committee of McGill University under protocol MCGL-5088. Animals were maintained in a pathogen-free environment and under standard housing conditions (23 ± 2°C in a 12 h light/12 h dark cycle) with ad libitum access to food and water. MRP8-Cre-ires/GFP (MRP8-Cre) transgenic mice were obtained from the Jackson laboratory (Bar Harbor, ME, USA; JAX, stock #021614). These mice express Cre recombinase under the control of the MRP8 promoter, which is active in myeloid cells.

### 2.2. Neutrophil-deletion of *F13a1* using MRP8-Cre driver mouse

For the development of neutrophil specific *F13a1* knockouts (*F13a1*^-/-MRP8^), MRP8-Cre mice were bred to *F13a1* flx/flx mice, which carry loxP sites inserted into exon 8 of *F13a1* gene, to delete *F13a1* gene in neutrophils. The knockout mice were selected based on the genotyping of pups using Selleck Direct PCR Kit and primers specific to LoxP (Forward: tctgggccaaaccaagtacctgg, Reverse: caagaccagactgtgcaaaggg) and Cre (Forward: gcggtctggcagtaaaaactatc, Reverse: gtgaaacagcattgctgtcactt) alleles. Qualitative RT-PCR using *F13a1* specific primers (Forward: cagttcgaagacggcatcct, Reverse: aacaagatcactgttgacctct) was also conducted to further confirm the knockout process.

### 2.3. Dietary intervention and weekly weight monitoring

All mice MRP8-Cre, *F13a1* flx/flx, *F13a1*^-/-MRP8^ were placed on specific diets at age of 4 weeks for 16 weeks. The diets including control diet (CD)(TD.08806) with 10% calorie from fat and high fat diet (HFD)(TD.06414) containing 60% calorie from fat were purchased from Envigo, headquartered in Indianapolis, Indiana, USA. Mice were weighed on a weekly basis to monitor the effect of specific diet on each group’s weight. *F13a1* flx/flx acted here as positive control with expected normal weight gain on HFD. The weight gain data was recorded and analyzed using GraphPad Prism v9. For better visualization, graphs were generated to represent weekly weight gain over 16 weeks on diets. The area under the curve (AUC) values were calculated using GraphPad Prism v9. The higher AUC represents higher weight gain.

### 2.4. Tissue collection, RNA extraction and cDNA synthesis

After 16 weeks on diet, the mice were euthanized by isoflurane followed by CO_2_ asphyxiation. The nose to tail of mice was measured as a parameter for growth. The white adipose tissue (WAT) depots, including visceral AT (VAT; epididymal) and subcutaneous AT (SAT; inguinal) were dissected, snap-frozen in liquid nitrogen, and stored at -80°C. Total RNA was extracted from snap-frozen tissues using the RNeasy Mini Kit (Qiagen, Germany), following the manufacturer’s protocol. Briefly, tissues were homogenized in QIAzol buffer, and RNA was purified using silica-membrane spin columns. The quantity and quality of RNA were assessed using a spectrophotometer, and the complementary DNA (cDNA) was synthesized from 1000ng of total RNA using the High-Capacity cDNA Reverse Transcription Kit (Applied Biosystems, USA). The reaction setup included the following components: 10x RT buffer (2.0 μL), 25x dNTP mix (100 mM, 0.8 μL), 10x RT random primers (2.0 μL), MultiScribe Reverse Transcriptase (1.0 μL), and nuclease-free water (4.2 μL). The reaction conditions were 25°C for 10 minutes, 37°C for 120 minutes, followed by a termination step at 85°C for 5 minutes.

### 2.5. Quantitative Real-Time PCR (qRT-PCR)

Each 20 µL reaction contained 9 µL (50 ng) synthesized cDNA, 10 µL TaqMan Fast Advanced Master Mix, and 1 μL of each TaqMan Gene Expression Assay. Relative gene expression of *Serpine 1* (Mm01204470_m1), *Ap1s1* (Mm00475917_m1) and *Adgre1*(Mm00802529_m1) was calculated using the 2^^(-ΔΔCt)^ method with TATA-box binding protein (*TBP*) (Mm01277044_m1) as the internal control for normalization.

### 2.6. Statistical analysis

Data were analyzed using GraphPad Prism software (version 9). Results are expressed as the mean ± SEM (standard error of the mean). Each experiment was performed in triplicate, with n representing the number of mice (*n* = 7). For comparisons involving more than two groups, one-way or two-way ANOVA was performed, as appropriate, with a Tukey post hoc test for multiple comparisons. The area under the curve (AUC) for weight gain was calculated with baseline 4-week weight set at zero. Differences were considered statistically significant at p-values < 0.05. Plots of the data were generated to visually assess trends and variability.

## 3. Results

### 3.1. MRP8-Cre and *F13a1*^-/-MRP8^ mice are resistant to weight gain on HFD

Diet-induced obesity models were created from MRP8-Cre mice, *F13a1* flx/flx, and *F13a1*^-/-MRP8^ models. *F13a1* flx/flx was used as a positive control with expected normal weight gain on HFD. The mice at age of 4 weeks, a starting point for diet challenge, showed similar weight (**Supplemental Figure 1**). Weekly weight monitoring showed significant difference in weight gain on HFD onwards of 8 weeks and resistance to weight gain in both MRP8-Cre and *F13a1*^-/-MRP8^ mice compared to the *F13a1* flx/flx control mice (**Figure 1A, B**). This difference persisted until the endpoint at 16 weeks and mice were visibly slimmer and smaller (**Figure 1C, D**). A similar pattern was observed in female groups, with differences showing after 10 weeks on HFD (data not shown). To determine if the weight difference was due to altered general growth and body size, nose-to-tail length was measured. The data indicated no significant differences in this parameter, suggesting normal general growth in all mouse models (**Figure 1E**).

**Figure 1:**
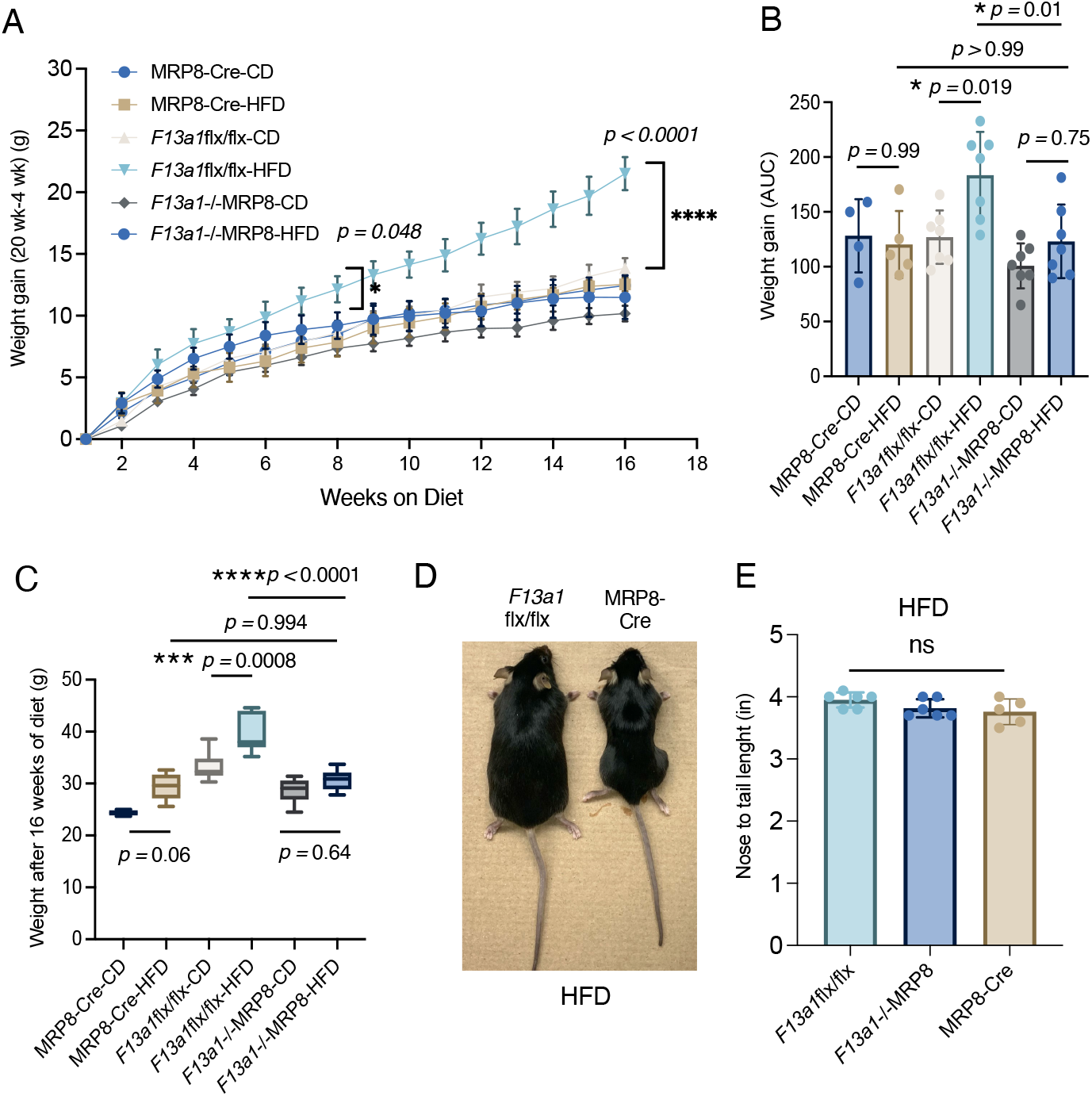
MRP8-Cre driver mouse does not gain weight on high-fat diet (HFD). **A. B**. Weight monitoring of MRP8-Cre driver mouse on HFD in comparison to *F13a1*flx/flx and *F13a1*-/-^MRP8^ (conditional knockout generated using MRP8-Cre model). Weekly weight gain and AUC of the weight gain curve are presented. The significant differences in weight gain begun to emerge after 8 weeks on HFD. **C**. Weights at the end point after 16-weeks on diet. MRP8-Cre sustained same weight on HFD as on CD whereas *F13a1*flx/flx (representing WT control genotype) gained significant weight on HFD. *F13a1*-/-^MRP8^ showed same resistance to weight gain on HFD as MRP-Cre on HFD. **D**. Photographs of *F13a1*flx/flx and MRP8-Cre after HFD feeding. MRP8-Cre appears slimmer. **E**. Nose-to-tail length measurements of *F13a1*flx/flx (control), *F13a1*-/-MRP8 and MRP8-Cre show no difference in general size of the mice. Data are presented as mean ± SD and significance was analyzed with two-way ANOVA. AUC data was normalized to weight at 4 weeks. Significance is presented as *p*-values and values <0.05 are considered significant (p< 0.05, *; p<0.001;**, p<0.001, *** and p<0.0001, ****).

### 3.2. WAT from MRP8-Cre mice show negligible *Serpine 1, as well as reduced Ap1s1* and *Adgre1* expression compared to *F13a1* flx/flx mice

To investigate whether the reported off-target genomic deletions of *Serpine1* and *Ap1s1* also resulted in deletions in AT depots, possibly affecting the weight gain, gene expression analysis of SAT (inguinal fat) and VAT (epididymal fat) was performed from the CD and HFD-fed mouse tissues. The analysis revealed that the expression of *Serpine1* in both AT depots and after both diets was indeed, dramatically and significantly down in MRP8-Cre and *F13a1*^-/-MRP8^ mice but increased in control model on HFD. Similar pattern was also observed for the expression of *Ap1s1* in VAT and SAT (**Figure 2A, B**). The MRP8-Cre background also generated SAT and VAT with significantly decreased macrophage infiltration (*Adgre1* - F4/80) suggesting protection from inflammation (**Figure 2A, B**).

**Figure 2:**
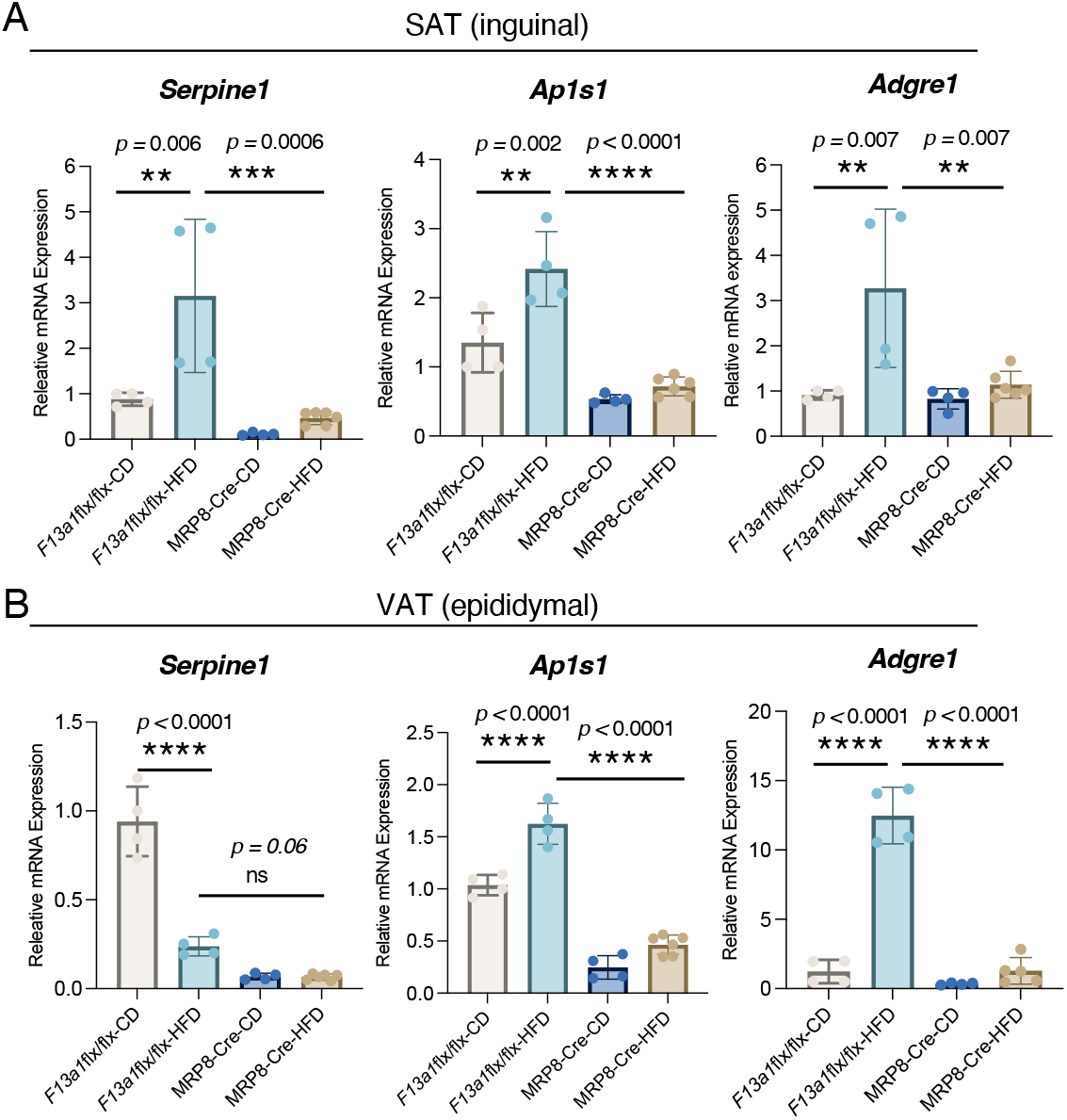
White adipose tissue depots of MRP8-Cre driver mouse are knockout for *Serpine1* and *Ap1s1* and show protection to macrophage infiltration on HFD. Gene expression analysis of *Serpine1* (PAI-1) *and Ap1s1* (AP1S1) and *Adgre1* (F4/80) in **A**. subcutaneous (inguinal) adipose tissue (SAT) and **B**. visceral (epididymal) adipose tissue VAT of MRP8-Cre mice (male) after 16 weeks on control diet (CD) and high-fat diet (HFD). HFD is expected to induce macrophage infiltration as observed in *F13a1* flx/flx on HFD (corresponding to a WT mouse genotype and positive control for diet-induced obesity model.

## 4. Discussion

PAI-1 inhibits fibrin lysis as part of hemostatic process to sustain blood clot. Recent advances have linked its function also to metabolic processes, AT inflammation, adipogenesis, development of insulin resistance, and glucose metabolism disturbances [15-17, 39, 40]. Emerging evidence shows that PAI-1 inhibition or genetic deletion has a positive effect on adipose tissue health and metabolic functions [14, 18, 41] as pharmacological inhibition of PAI-1 using PAI-039 or complete deletion of the PAI-1 gene lead to a significant decrease in macrophage infiltration into AT and improved metabolic status in HFD-induced models[14]. In addition, targeting PAI-1 with chemical inhibitors ameliorates AT inflammation, atherosclerosis and associated metabolic dysfunctions[42]. Mechanistically, PAI-1’s role to weight gain in mice was linked to enhanced hypothalamic leptin resistance but roles directly in AT are also likely based on recent advances in our understanding of the essential roles of fibrinolytic system on AT health and inflammation[18]

[12]. Factor XIII-A, transglutaminase enzyme that stabilizes fibrin clot was originally linked to obesity as putative causative gene in a GWAS study on obesity-discordant monozygotic twins[43], however, global deletion of FXIII-A in mice does not contribute to weight gain albeit the mice have significantly improved insulin sensitivity on HFD compared to controls [13]. Our recent transcriptome-wide association studies (TWAS) on adipocytes and adipose tissues from monozygotic twins linked the *F13A1* gene to neutrophil activation justifying the question if this FXIII-A in neutrophils is relevant to AT health [44-46]. Generation of neutrophil *F13a1* knockout model using MRP8-Cre mice revealed that MRP8-Cre background itself creates resistance to weight gain, a finding confirmed in this paper using the MRP8-Cre alone. It is likely that the recently reported off-target effects in the MRP8-Cre [37] are the underlining cause of the weight phenotype. A recent study [38] also showed the impact of MRP8-Cre transgene on innate immunity, probably due to the interfering with expression of genes near to the insertion site.

Our gene expression analysis verified the complete deletion of *Serpine1* in SAT and VAT along with a significant reduction in *Ap1s1* expression. AP1S1, a member of the adaptor protein complex family, contributes to cargo transport between intracellular organelles [47-49]. To date, there is no data available on the protein’s contribution to weight modulation and metabolic health.

So far, MRP8-cre mice have been used to generate several neutrophil specific knockouts for analysis of metabolic effects. These involve PAD4^-/-MRP8^, CXCR2^-/- MRP8^, CXCR4^-/- MRP8^, and α9^-/- MRP8^ [50-53]. MRP8-Cre model was not used as control. PAD4^-/- MRP8^ study showed a reduction in weight gain from 7 weeks onwards of HFD, and showed no increase in cardiac collagen, and exhibited a decreased thrombus formation after 10 weeks of metabolic challenge [50]. Additionally, the obese α9^-/-MRP8^ model showed improved stroke outcomes and reduced thrombosis susceptibility [53]. A study on the effects of neutrophil aging on metabolic profiles in CXCR2^-/- MRP8^ mice, challenge with a 20-week HFD, revealed protection from metabolic alterations and reduced proinflammatory macrophage recruitment in AT. These mice also exhibited resistance to weight gain [52]. While our report is based on simple weight gain observations, it is important to emphasize that reduced weight gain on HFD will naturally improve many metabolic outcomes, including insulin sensitivity, glucose clearance and AT inflammation. Furthermore, global elimination of PAI-1 in the MRP8-Cre may cause number of other phenotypes, particularly those related to thrombosis and thromboinflammation. We thus encourage re-investigating some of the discovered effects using the MRP8-Cre as control. Our report underscores the critical need for thorough characterization of genetically engineered models and the use of appropriate mouse controls in all studies.

## 5. Conclusion

In conclusion, our study underscores the confounding effects of MRP8-Cre mice when investigating weight gain and metabolic parameters and emphasizes the importance of including appropriate controls, for any new genetic models.

## Acknowledgments

This research was funded by the Canadian Institutes of Health Research (CIHR), with grants awarded to Mari T. Kaartinen (PJT153089). Montreal, Canada. Fatemeh Soltani was supported by stipends from the Faculty of Medicine and Health Sciences of McGill University, and by a scholarship from Le Fonds de recherche du Québec - Santé (FRQ-S).

## CRediT authorship contribution statement

**Fatemeh Soltani:** Conceptualization, Investigation, Methodology, Writing – original draft and review & editing. **Cedric Duval**; Methodology, Writing – review & editing. **Robert A. S. Ariëns:** Methodology, Writing – review & editing. **Mari T. Kaartinen:** Conceptualization, Funding, Project administration, Supervision, Visualization, Writing– review & editing.

## Animal ethics committee approval

The animal study protocol was approved by the Animal Care Committee of McGill University (protocol code MCGL-5188, May 3, 2023).

## Figure Captions

**Supplemental Figure 1: Weights of mice at 4-week age prior to starting special diets**. No statistical differences were detected.

