## Supplemental Figure 1 for "Off-target genomic effects in MRP8-Cre driver mice complicate its use in weight gain and metabolic studies"

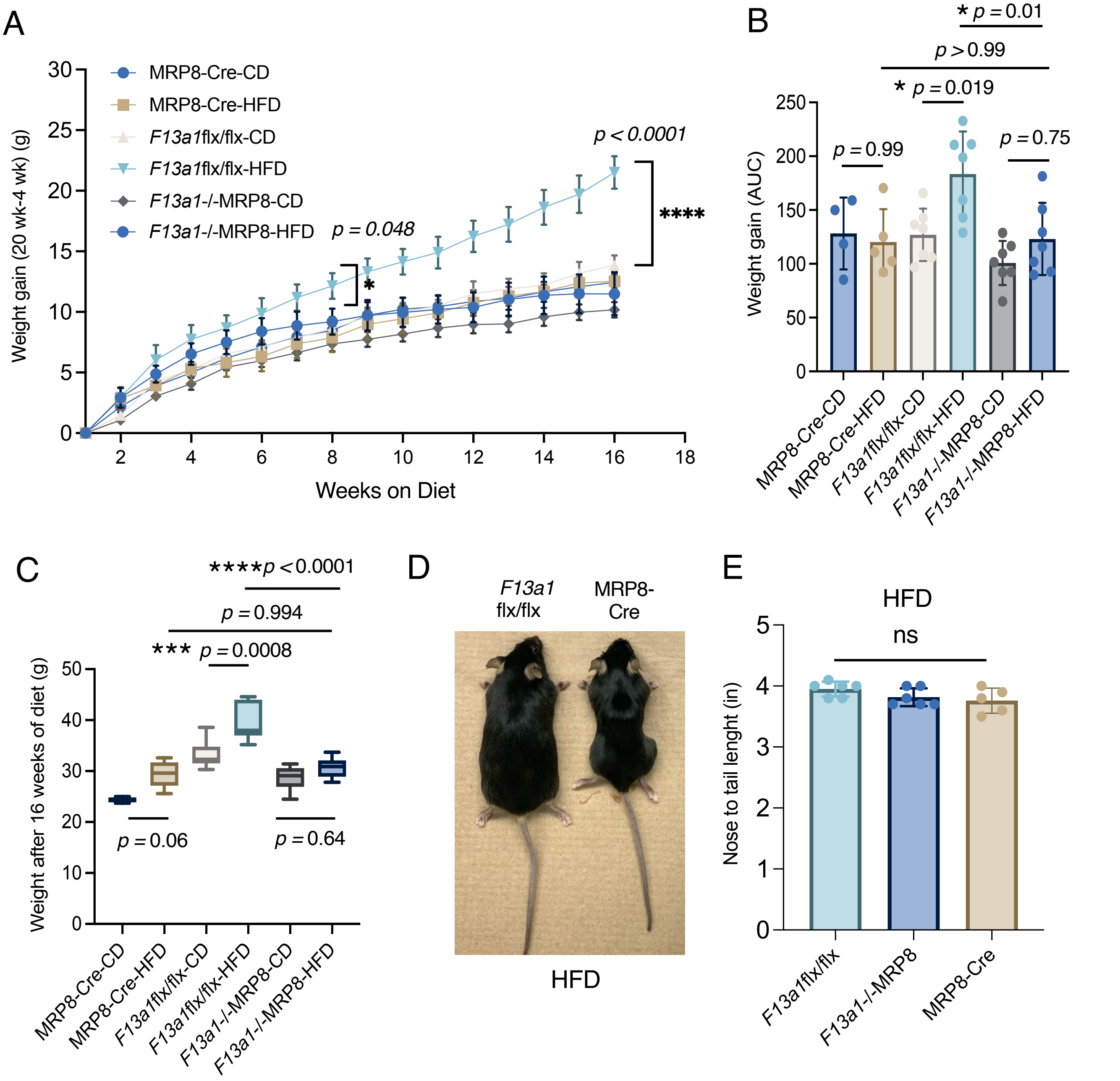


**Figure 1: MRP8-Cre driver mouse does not gain weight on high-fat diet (HFD). A. B.** Weight monitoring of MRP8-Cre driver mouse on HFD in comparison to *F13a1*flx/flx and *F13a1*-/-$^{\mathrm{MRP8}}$ (conditional knockout generated using MRP8-Cre model). Weekly weight gain and AUC of the weight gain curve are presented. The significant differences in weight gain begun to emerge after 8 weeks on HFD. **C.** Weights at the end point after 16-weeks on diet. MRP8-Cre sustained same weight on HFD as on CD whereas *F13a1*flx/flx (representing WT control genotype) gained significant weight on HFD. *F13a1*-/-$^{\mathrm{MRP8}}$ showed same resistance to weight gain on HFD as MRP-Cre on HFD. **D.** Photographs of *F13a1*flx/flx and MRP8-Cre after HFD feeding. MRP8-Cre appears slimmer. **E.** Nose-to-tail length measurements of *F13a1*flx/flx (control), *F13a1*-/-MRP8 and MRP8-Cre show no difference in general size of the mice. Data are presented as mean ± SD and significance was analyzed with two-way ANOVA. AUC data was normalized to weight at 4 weeks. Significance is presented as *p*-values and values <0.05 are considered significant (p< 0.05, *; p<0.001;**, p<0.001, *** and p<0.0001, **** ).

### Figure 2

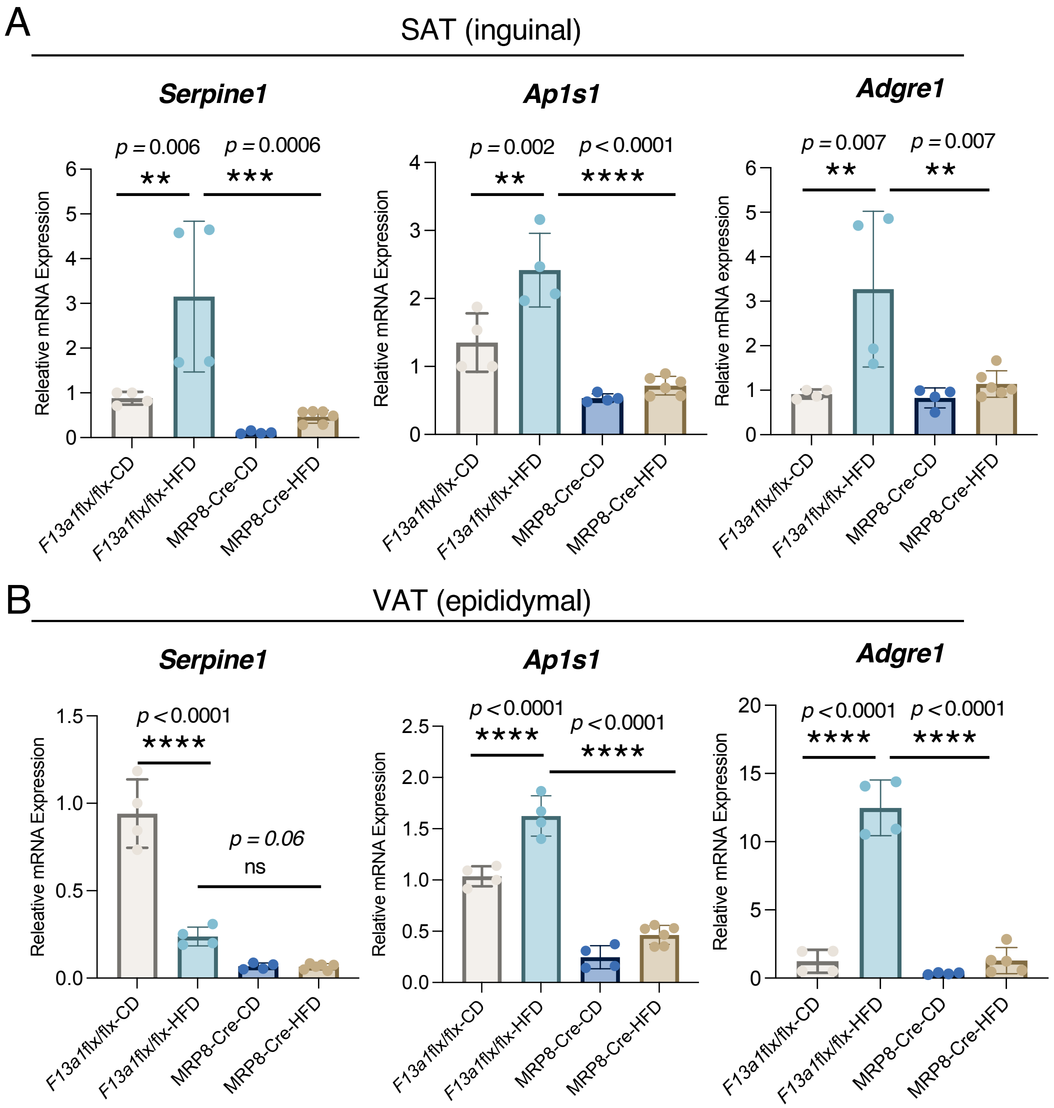


**Figure 2: White adipose tissue depots of MRP8-Cre driver mouse are knockout for *Serpine1* and *Ap1s1* and show protection to macrophage infiltration on HFD.** Gene expression analysis of *Serpine1* (PAI-1) *and Ap1s1* (AP1S1) and *Adgre1* (F4/80) in **A.** subcutaneous (inguinal) adipose tissue (SAT) and **B.** visceral (epididymal) adipose tissue VAT of MRP8-Cre mice (male) after 16 weeks on control diet (CD) and high-fat diet (HFD). HFD is expected to induce macrophage infiltration as observed in *F13a1* flx/flx on HFD (corresponding to a WT mouse genotype and positive control for diet-induced obesity model.

Supplementary Figure 1

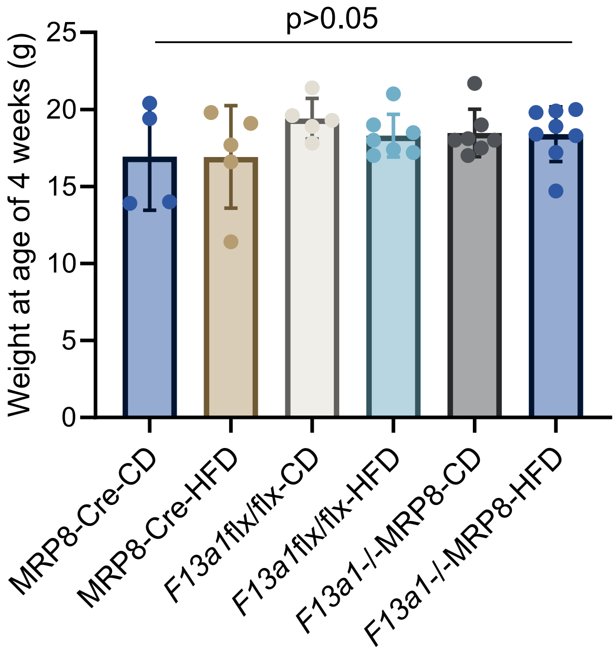


**Supplemental Figure 1: Weights of mice at 4-week age prior to starting special diets.** No statistical differences were detected.
